# Real-Time Segmentation and Classification of Birdsong Syllables for Learning Experiments

**DOI:** 10.64898/2025.12.19.695629

**Authors:** Nils Riekers, Jacqueline Laura Göbl, Franziska Heubach, Lena Veit

**Affiliations:** Neurobiology of Vocal Communication,Institute for Neurobiology, University of Tübingen, Germany

## Abstract

Songbirds are essential animal models for studying neuronal and behavioral mechanisms of learned vocalizations. Bengalese finch (*Lonchura striata domestica*) songs contain a limited number of acoustically distinct syllable types, which are combined into variable sequences. This makes them ideal to investigate the composition of vocal sequences. Many closed-loop experiments require the online recognition of a specific target syllable while the bird is singing, for example when introducing specific song modifications through reinforcement learning. In this protocol, a specific target syllable is covered with a short burst of white noise, which masks auditory feedback of the bird’s own song, and leads the bird to avoid the targeted syllable in future song renditions. Existing tools for this learning protocol require manual creation of spectral templates and have limited flexibility to adapt to new experiments. We here present Moove (Marking Online using Only the Onsets of Vocal Elements), a novel approach to real-time syllable segmentation and classification of Bengalese finch songs. A convolution-based audio encoder with a simple multi-layer perceptron is used for onset and offset segmentation of individual syllables, and syllable classification is performed by a convolutional neural network. Crucially Moove classifies syllables using only acoustic information from the first part of the syllable, while the syllable is still ongoing. This allows reinforcement or punishment to be applied with high accuracy and short latency after syllable onset, enabling effective operant conditioning experiments. The fast online annotation of all recorded syllables allows targeting syllables based on their sequence context, while keeping classification itself free from influence by acoustic information from surrounding syllables. To validate this tool, we conducted a learning experiment on one adult male Bengalese finch. The bird learned to avoid the targeted syllable sequence with comparable outcomes to previous learning experiments. Our results show that Moove can correctly segment and classify Bengalese finch syllables in real time for learning experiments. Moove could be used for other closed-loop experiments such as manipulation of auditory feedback, song-triggered neuronal microstimulation, optogenetics, or reward delivery. This makes Moove a crucial tool for future investigations of birdsong sequencing.

## Introduction

Songbirds are ideally suited to study learned vocal communication, since they learn their song by vocal imitation of conspecifics, in a process that shares parallels with human speech learning (Brainard & Doupe, 2013; Doupe & Kuhl, 1999). In many species, song is composed of acoustically discrete vocal elements, called syllables. Bengalese finches flexibly arrange these syllables into variable sequences following learned syntactic rules (Koparkar et al., 2024; Okanoya, 2004; Okanoya & Yamaguchi, 1997; Veit et al., 2021). We here use letters (e.g., ‘a’, ‘b’, ‘c’) to label distinct syllable types. Syllable sequences in Bengalese finch song contain meta-level structures such as chunks, where syllables follow a fixed order (such as ‘a-b-c’), branch points, where multiple different syllables may follow the same syllable type (such as ‘c-d’ 40% of instances and ‘c-e’ 60% of instances) as well as repeat phrases, where the same syllable type is repeated a variable number of times (such as ‘d-d-d-d-d’ repeated between 3-8 times in each instance of the phrase) (Koparkar et al., 2024; Seki et al., 2008; Suge & Okanoya, 2010). Studying the mechanisms underlying the sequential composition of birdsong can help to reveal the neuronal basis for syntactic ordering of actions in motor skills.

Immense progress has been made deciphering the neuronal mechanisms responsible for song production and learning by interfering with adult song in closed-loop experiments (Andalman & Fee, 2009; Sober & Brainard, 2012; Tumer & Brainard, 2007). These experiments require the online recognition of target syllables and immediate delivery of feedback while the bird is singing. For example, important experimental approaches include song-triggered manipulations of auditory feedback (Gadagkar et al., 2016; Riedner & Adam, 2020; Sakata & Brainard, 2006; Sober & Brainard, 2009; Tumer & Brainard, 2007), visual feedback (Cynx, 1990; Franz & Goller, 2002; Seki et al., 2008; Zai et al., 2020) or somatosensory feedback (McGregor et al., 2022), song-triggered optogenetic stimulation (Hisey et al., 2018; Roberts et al., 2012; Xiao et al., 2018), microstimulation (Ashmore et al., 2005; Kao et al., 2005; Moll et al., 2023), or other behavioral contingencies such as delivery of video playback (Carouso-Peck & Goldstein, 2019; Kawaji et al., 2024). One key protocol we especially tried to adapt is introducing specific learned changes to the song using reinforcement learning. In these protocols, a specific target syllable is masked by a short burst of white noise (WN). Since the WN disturbs auditory feedback of the bird’s own song, the bird will typically introduce song modifications that avoid triggering the WN playback on future renditions. These changes are learned modifications of the song which occur in subsequent song bouts, even in the absence of WN. This protocol has been used to modify syllable pitch (Andalman & Fee, 2009; Canopoli et al., 2014; Tian et al., 2023; Tian & Brainard, 2017; Tumer & Brainard, 2007; Warren et al., 2012; Zai et al., 2024), timing (Ali et al., 2013; Tachibana et al., 2017) as well as song sequencing in Bengalese finches (Fortkord & Veit, 2025; Kawaji et al., 2024; Veit et al., 2021; Warren et al., 2012).

Syllable annotation is usually performed in a two-stage process. Syllable segmentation refers to detecting the onset and offset of individual vocal elements within a continuous song recording, while syllable classification assigns each detected element to a specific syllable type based on its acoustic features. Current solutions for closed-loop experiments range from manual triggering by humans (Carouso-Peck & Goldstein, 2019; Phaniraj et al., 2022) to spectral template-matching (Fortkord & Veit, 2025; Kao et al., 2005; Tumer & Brainard, 2007) to neural network solutions (Kawaji et al., 2024; Steinfath et al., 2021). Template-based solutions like EvTAF, a LabView program, skip the syllable segmentation step and match templates on a continuous input stream. They combine advantages such as low latency and simple setup, but they require manually designed templates for target syllables, which need to be manually adapted to syllable changes introduced by the bird. The creation of an optimal template is a challenging process and substantially influences the accuracy of syllable recognition (Kulkarni & Troyer, 2020). EvTAF can recognize target syllables in specific sequences by combining multiple templates with Boolean logic and set latencies between template matches, but this quickly leads to increasingly complex template combinations for birds with variable sequencing. Therefore, EvTAF can be cumbersome to use and adapting it to new experiments requires LabView programming skills. Recently, neural network-based solutions have been introduced that are capable of online annotation of birdsong syllables (Kawaji et al., 2024; Schulthess et al., 2023; Steinfath et al., 2021). While these approaches represent a major advance and offer high classification accuracy without the need for manual template design, all currently available approaches classify syllables only when syllable detection is completed, after syllable offset. This introduces latencies which are much longer than the fast closed-loop capability of EvTAF. For such experiments, it can be essential to target syllables while they are still being vocalized (Charlesworth et al., 2011; Tumer & Brainard, 2007). Therefore, there is no readily available solution to reliably target syllables before their offset with modern neural network approaches.

This motivated us to develop Moove (Marking Online using Only the Onsets of Vocal Elements) as a novel approach to real-time birdsong annotation. An important goal was to replace and extend the functionalities of EvTAF, while at the same time ensuring compatibility through similar file types and annotation formats to facilitate adoption by research groups which are already using this software (including our own). Further goals were to use a Python-based system, which is easier to modify for most neurobiology students, to work without specialized and expensive hardware, and to adapt real-time targeting to more variable song amplitude levels, which we experience in relatively large experimental cages where the bird can have varying distances from the microphone. Figure 1 illustrates the functionality of Moove, which performs segmentation and classification while the syllable is still being sung, allowing for fast feedback delivery.

**Figure 1.**
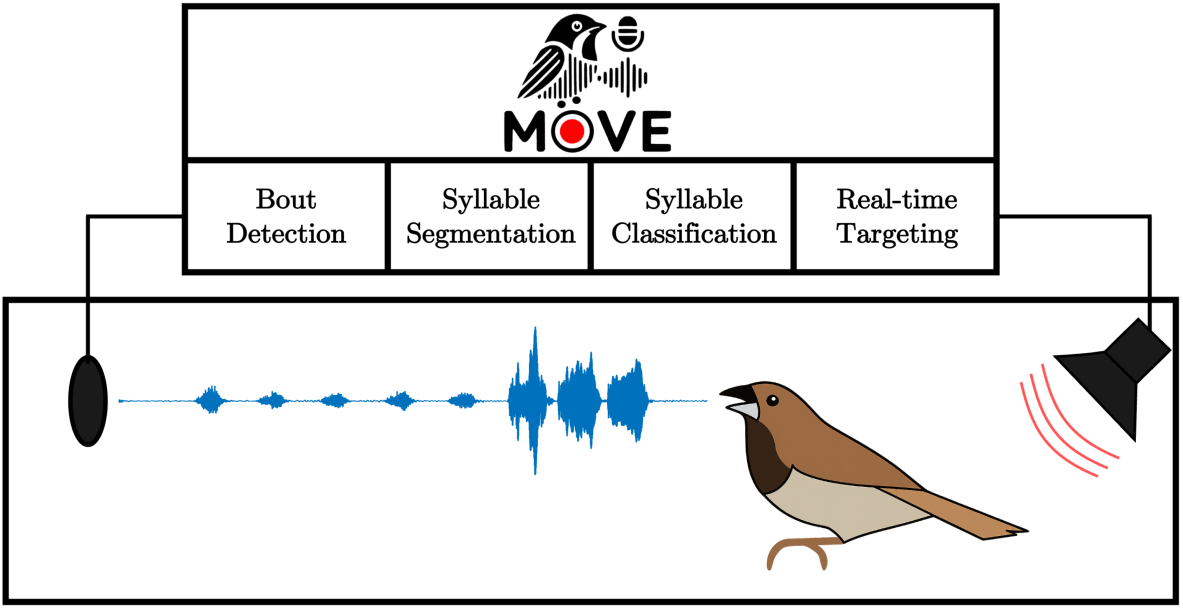
Functional overview of Moove. The system continuously monitors vocalizations from a singing Bengalese finch and performs four sequential processing steps: (1) Bout Detection identifies song bouts from continuous audio input, (2) Syllable Segmentation detects onsets and offsets of individual syllables within detected bouts, (3) Syllable Classification assigns syllable types using only the initial portion of each syllable, and (4) Real-time Targeting triggers feedback stimuli (e.g., white noise playback) while the target syllable is still being produced. The diagram illustrates the complete closed-loop system from audio acquisition (microphone, left) through neural network processing (Moove pipeline, top) to feedback delivery (speaker, right), enabling operant conditioning experiments with short latencies.

## Methods

Moove (Marking Online using Only the Onsets of Vocal Elements) is a Python package for real-time birdsong syllable segmentation and classification using neural networks. The package consists of MooveTAF (Moove Targeted Auditory Feedback) for song recording and real-time targeting, and MooveGUI for data preprocessing, manual labeling, and network training. The source code is publicly available at https://github.com/veitlab/moove.

### MooveTAF: Recording and real-time targeting

MooveTAF was developed to provide audio recording and real-time targeting of birdsong syllables, enabling online segmentation, classification, and feedback triggering. It uses the Python package *sounddevice* (Geier et al., 2020) as an interface to PortAudio (Bencina & Burk, 2001), and supports multiple native audio APIs. The name and several of its features are inspired by the LabView program EvTAF (Tumer & Brainard, 2007).

For real-time segmentation and classification of song syllables, a key design requirement was to balance high accuracy with fast inference time. The inference speed of the system must be faster than the rate at which new audio chunks, small consecutive sections of the audio stream, are processed, to prevent delays from accumulating and negatively impacting the tool’s real-time capability. The size of audio chunks is therefore one of the major contributors to latency in real-time audio classification (Steinfath et al., 2021). Smaller chunks reduce latency but require faster inference and limit the temporal context available for classification. To address this, a two-stage architecture was implemented that operates on very small chunk sizes, allowing rapid syllable onset detection and thereby reducing latency.

### Architecture of MooveTAF

Our solution consists of one dedicated network for the detection of syllable segments in a continuous stream of audio data (the segmentation network) and a second network to classify these syllable segments (the classification network). After the segmentation network has detected a syllable onset, the following audio data is collected over a short period of time and then transferred to the classification network. This architecture is similar to the one utilized by Schulthess et al. (2023), which also has a two-stage architecture using a dedicated segmentation and classification stage.

The segmentation network (Fig. 2A) determines whether each incoming audio chunk is part of a syllable segment. It consists of a convolutional encoder to extract features and a multi-layer perceptron (MLP) to determine segment membership (syllable/gap). Using the default parameters, this network makes approximately 690 inferences per second with a chunk size of 64 samples at 44.1 kHz. Due to this high number of inferences, errors inevitably occur even at high accuracy levels. Therefore, a sliding window algorithm (Fig. 2B) was implemented which identifies syllable onsets when a pre-defined number of chunks within a sliding window are classified as part of a syllable segment (default: 3 out of 5 consecutive chunks). This post-processing step reduces false positives from individual chunk predictions and yields continuous syllable segments that correspond well to syllables in the spectrogram (Fig. 2C).

**Figure 2.**
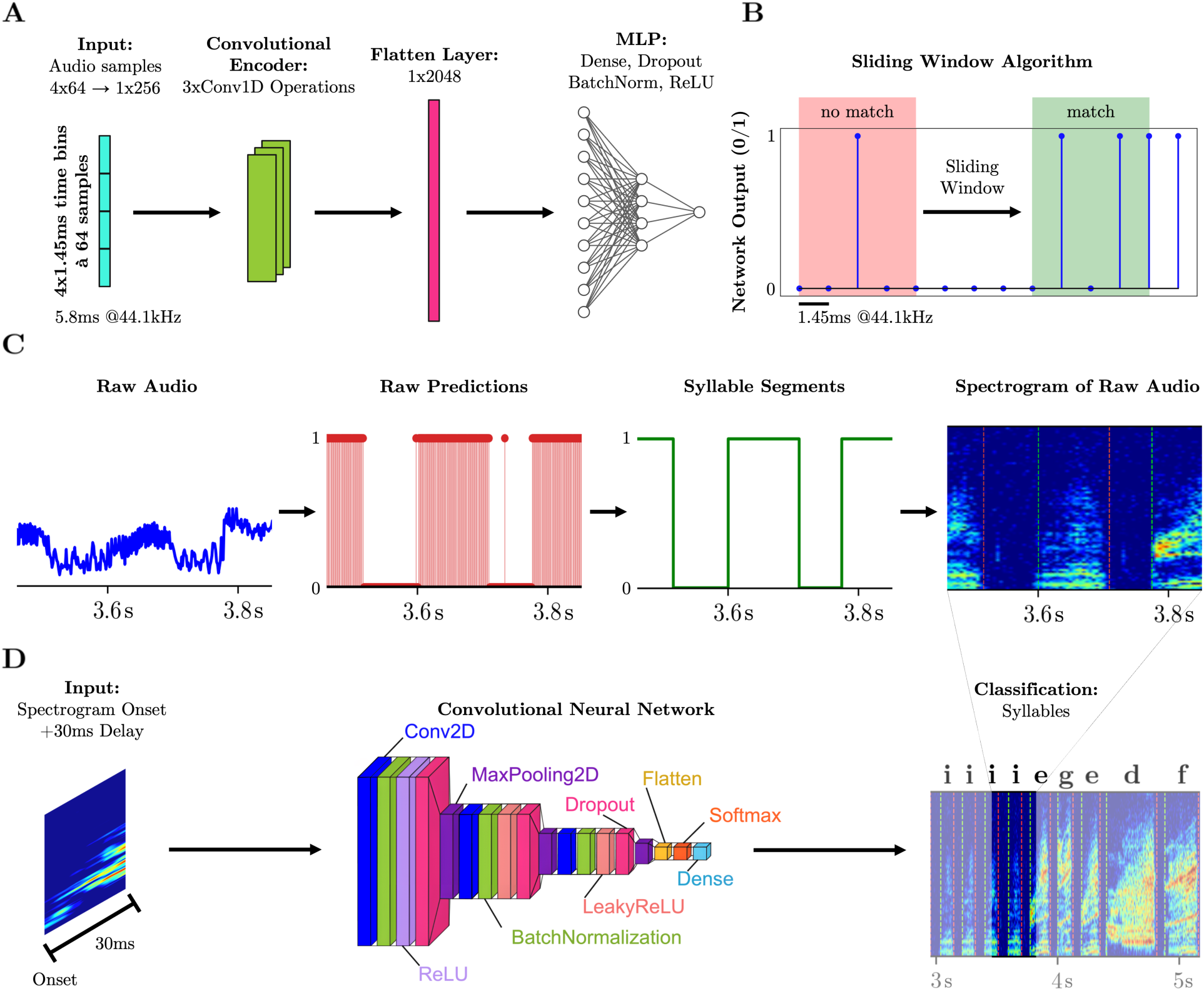
Architecture of the two-stage neural network system in MooveTAF. (A) Segmentation network architecture. The network receives consecutive audio chunks (64 samples at 44.1 kHz) together with the three preceding chunks as input. A convolutional encoder extracts features from the raw audio, followed by a multi-layer perceptron (MLP) that predicts whether the current chunk belongs to a syllable segment (output: 0 or 1). (B) Sliding window algorithm for onset detection. A syllable onset is confirmed when at least 3 out of 5 consecutive chunks in a sliding window are classified as belonging to a syllable segment (green). When fewer than 3 chunks are classified as syllable (red), no onset is detected. (C) Example of segmentation output showing the complete processing pipeline. From left to right: Raw audio waveform, raw binary predictions from the segmentation network (red bars with white vertical lines indicating individual chunk predictions), merged syllable segments after applying the sliding window algorithm (green boxes), and spectrogram of the raw audio with vertical dashed lines marking the detected syllable boundaries. (D) Classification network architecture. After syllable onset detection, 30 ms of audio are buffered and converted to a spectrogram. This spectrogram is processed by a convolutional neural network (CNN) consisting of three convolutional blocks (Conv2D, BatchNormalization, LeakyReLU/ReLU, Dropout, MaxPooling2D), followed by a flatten layer, a fully connected layer, and softmax activation to assign a syllable label (e.g., ‘a’, ‘b’, ‘c’).

Once a syllable onset is detected, audio data are buffered for a short period (default: 30 ms; ∼30.47 ms at 44.1 kHz, chunk size 64) and transformed into a spectrogram, which serves as input for the classification network (Fig. 2D). The classification network, a CNN, then assigns a label to the syllable segment based on this input. If the classified syllable sequence matches a user-defined target sequence, MooveTAF can trigger a feedback stimulus (e.g., white noise playback), enabling real-time closed-loop experiments.

For real-time targeting, one or several target sequences can be defined by the user. Once targeting is enabled, MooveTAF continuously monitors the detected syllable stream and applies regular expression-based pattern matching to determine whether the current sequence matches a predefined target. This approach provides substantial flexibility to accommodate song variability and to account for occasional classification errors. For example, the pattern `[jc]a$` triggers feedback for sequences ending in either *‘*j-a*’* or *‘*c-a*’*. Regular expression targeting rules can include or exclude specific sequence contexts or specify a range of repeat counts for repeat phrases. This enables precise experimental control over which vocal sequences trigger feedback.

If the currently classified syllable sequence matches the target pattern, a feedback stimulus is triggered. A catch-trial rate can be set to omit triggering in a defined proportion of bouts. By default, the feedback stimulus consists of playback of a user-provided wav file (e.g. white noise), though the system can be readily adapted for other feedback modalities. All parameters of the targeting behavior, such as the stimulus file and catch-trial rate, are user-definable in the config file. If multiple playback stimulus files are provided, MooveTAF randomly selects from the options with replacement. If multiple target sequences are defined, one is randomly selected at the beginning of each new song bout and remains active for its duration.

### MooveGUI: Data preprocessing, labeling and network training

Before syllable targeting can be performed, baseline recordings must be collected using MooveTAF in recording mode (segmentation, classification, and feedback disabled). Bouts are detected and saved via an amplitude threshold-based method using criteria such as minimum bout length. As illustrated in Figure 3, all subsequent data preprocessing and training steps are conducted in MooveGUI, which is fully integrated into the Moove Python package. MooveGUI provides a user-friendly, Tkinter-based graphical interface that allows users to execute all necessary preprocessing steps interactively and to train both neural networks directly within the tool. The segmentation and classification networks are trained separately for each bird, based on its individual song repertoire. Once training is complete, the resulting models can be loaded into MooveTAF to perform real-time segmentation, classification, and targeting during closed-loop experiments. This semi-automated pipeline thus provides a cohesive workflow from data collection to model deployment. The following sections describe each step of this preprocessing and training pipeline in detail.

**Figure 3.**
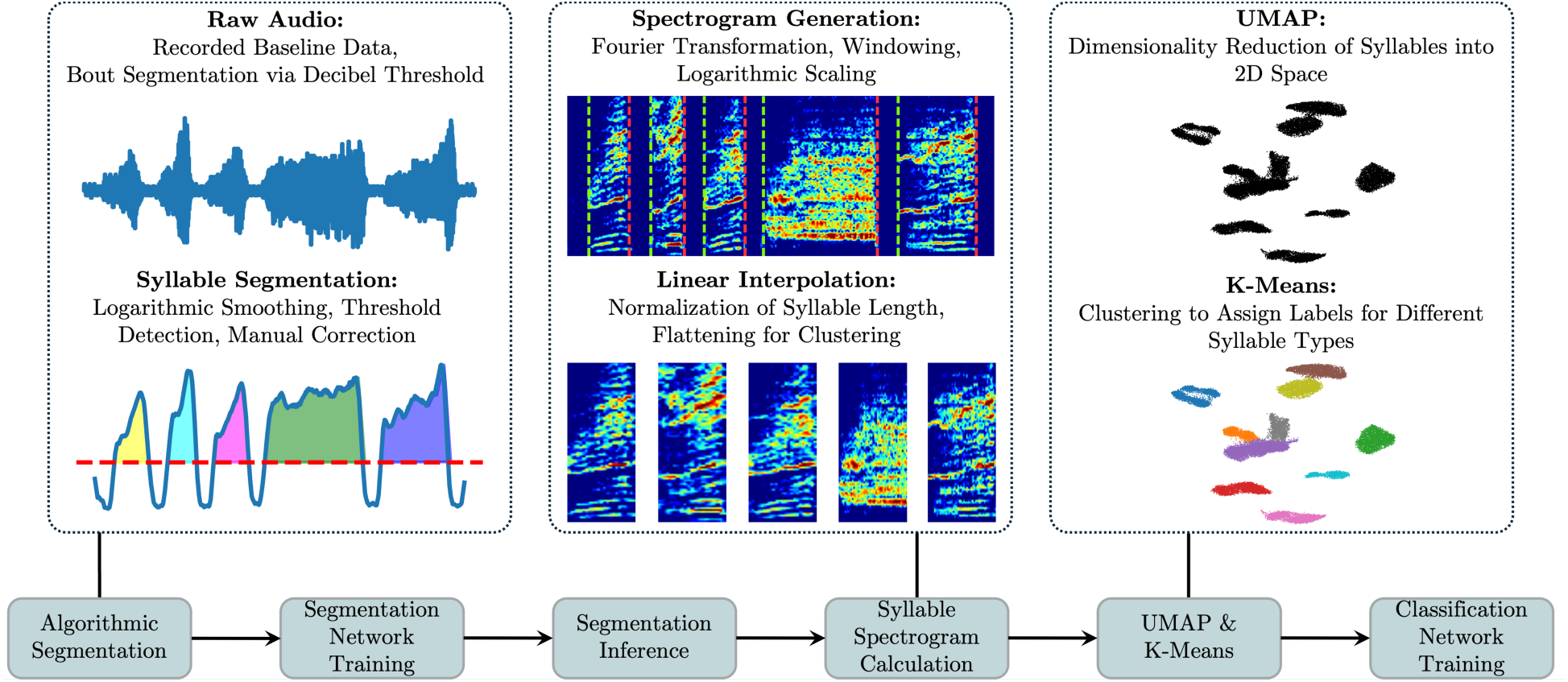
Data preprocessing and network training pipeline in MooveGUI. Left panel: Syllables are first segmented using an algorithmic amplitude-threshold method and, if necessary, manually corrected. The verified segments are then used to train the segmentation network. Subsequently, the trained network is applied to re-segment the data, ensuring consistency with the segmentation that will be performed during real-time experiments. Middle panel: From the resulting syllable segments, spectrograms are computed using a Short-Time Fourier Transform (STFT), normalized by linear interpolation to unify syllable length, and flattened into feature vectors. Right panel: These vectors are projected into a two-dimensional space using UMAP to visualize acoustic similarity between syllables, and k-Means clustering is applied to obtain initial syllable labels (colored clusters). Cluster assignments can then be manually refined via the GUI in the UMAP space or in the spectrogram to correct mislabels or merge entire clusters. Finally, the classification network is trained on spectrogram inputs paired with the curated labels, using the same temporal window (e.g. 30 ms) as during later online inference.

#### Algorithmic Segmentation

In the first step, each bout of raw audio data is segmented based on an amplitude-based threshold, using the algorithm implemented in *evfuncs* (Nicholson, 2021), which applies bandpass filtering, amplitude squaring, smoothing, and conversion into decibel values. An amplitude threshold is then used to detect syllable onsets and offsets, while segments shorter than a minimum duration are excluded, and segments separated by silent intervals shorter than a minimum duration are merged. This functionality has been integrated into MooveGUI, which also allows manual refinement of onset and offset times. Since algorithm-driven segmentation is prone to over- and under-segmentation (Cohen et al., 2022), manual corrections are often required to obtain consistent segmentation and thus accurate training data for the segmentation network.

#### Segmentation Network Training

Once a sufficient number of training bouts have been segmented, the segmentation network can be trained. The training can proceed iteratively: the manually segmented bouts are used to train an initial model, which is then refined using bouts segmented by the model, which are manually corrected and added to the training data. This process allows large numbers of training bouts to be obtained relatively quickly, gradually improving the model with each iteration (Steinfath et al., 2021).

Segmented data are randomly divided into training, validation, and test subsets. Downsampling is applied by default to balance the number of samples per label. The segmentation task is treated as a binary classification problem (syllable chunk vs. non-syllable chunk) using binary cross-entropy with logits as the loss function. Training is performed with the Adam optimizer, and early stopping is applied to prevent overfitting when validation loss no longer improves.

#### Segmentation Inference

Once the segmentation network is trained, it is applied to re-segment the training data using the same sliding window algorithm that will be used during real-time operation. This re-segmentation ensures consistency between offline training data and online syllable detection.

#### Syllable Spectrogram Calculation

For each syllable segment, spectrograms are computed using a Short-Time Fourier Transform (STFT). The same STFT parameters (window length, overlap, FFT size, and frequency cutoffs) are applied to all syllables. Syllable durations are normalized by linear interpolation to obtain equal-size spectrograms (Koparkar et al., 2024), and the amplitude of each spectrogram is converted into decibel values. The resulting spectrograms serve as input for subsequent dimensionality reduction and clustering.

#### UMAP & K-Means

The spectrograms of detected syllable segments are projected into a two-dimensional space using the Uniform Manifold Approximation and Projection (UMAP) (McInnes et al., 2018) algorithm (Kawaji et al., 2024; Koparkar et al., 2024; Sainburg et al., 2020; Steinfath et al., 2021), where syllables with similar spectral structure cluster together, resulting in distinct clusters for different syllable types in Bengalese finch song. The k-Means algorithm (MacQueen, 1967) is then applied to these embeddings to assign each syllable segment to one of the clusters and thus apply labels. UMAP parameters (number of neighbors and minimum distance) and number of clusters for k-means can be adjusted by the user.

#### Manual Adjustments and Verification of Cluster Membership (GUI-based)

MooveGUI includes an interface for manual adjustment of cluster memberships, where users can visually inspect the assigned labels in the 2D UMAP projection and directly re-assign data points. This allows manual splitting or merging of clusters or correction of mislabels.

#### Classification Network Training

The classification network is trained on spectrograms of syllable segments together with their manually verified labels. To enable low-latency online classification, only a fixed temporal window after each detected onset (default: 30 ms) is used as input during both training and inference. The Classification Network is optimized for multi-class classification using cross-entropy as the loss function. To improve generalization, data augmentation techniques such as noise addition, frequency masking, and time masking are applied to a subset of training samples. Dataset splits, normalization, Adam optimization, and early stopping are performed as described for the segmentation network training. After training the classification network, Moove can be used to reliably distinguish syllable types online during real-time experiments.

### Animals

Five male Bengalese finches were used to test different features of Moove across multiple experiments. Birds were housed in sound-attenuating chambers (120 × 50 × 50 cm) under a 14:10 h light–dark cycle at 25 °C and 50 % humidity, with ad libitum access to food, water, and standard enrichment (sand baths, nests, fresh greens). Vocalizations were recorded using a Rode M5 MP microphone (Australia) connected to a Steinberg UR12/IXO12 audio interface (Germany), with a JBL Control 1 Pro loudspeaker (USA) used for playback. All recordings were made using Moove’s default configuration settings as specified in the documentation. All experimental procedures were approved by the Regierungspräsidium Tübingen (Germany) and conducted in compliance with animal welfare regulations.

### Experiment 1: System performance characterization

Using networks trained on each bird’s individual song repertoire, we characterized system latency and tested position-specific repeat targeting capabilities in two birds. To characterize latency, we targeted a fixed syllable sequence and measured the time interval between syllable onset detection and white noise playback (Fig. 6). We tested different audio buffer sizes (64, 128, and 512 samples) and mode settings using the Yamaha Steinberg USB Driver on Windows. For each configuration, we collected at least 50 triggered instances to obtain reliable latency distributions. To demonstrate the ability to target specific repetitions within repeat phrases, we targeted the fifth occurrence of syllable ‘a’ and delivered position-specific auditory feedback by playing back different auditory stimuli (syllables ‘c’ or ‘d’) for different instances (Fig. 7).

### Experiment 2: Sequence modification learning experiment

A Bengalese finch with a clear branch point in its song was used for the learning experiment. Five days of baseline song were recorded with real-time processing disabled. These data were used to train the segmentation and classification networks. The trained model was then used for four consecutive days of white noise training, targeting a specific syllable sequence to reduce its occurrence after the branch point. An 80 ms white noise stimulus was played to overlay the targeted syllable. Catch trials (10%) were included where no feedback was given. Two post-screening sessions conducted after training assessed the persistence of learning. The bird was housed in a group aviary between baseline, training, and post-screening sessions.

## Results

We developed Moove, a two-stage neural network system for real-time segmentation and classification of Bengalese finch syllables. The system achieves syllable classification during ongoing syllables by using only the first 30 ms of acoustic information after syllable onset.

### Segmentation performance

We tested segmentation performance on five Bengalese finches with different songs. Segmentation networks were trained separately for each bird to distinguish syllable segments from gaps in the audio stream. Training data ranged from 108 to 576 manually verified bouts across birds (mean: 234 ± 195 bouts), which were randomly split into training (70%), validation (15%), and test (15%) subsets (see Table S1 for individual bird data). Across all birds, training and validation accuracies converged to high levels (Fig. 4A), with test accuracies ranging from 94.30% to 98.50% (mean: 95.91 ± 1.70%). Correspondingly, training and validation losses decreased to low levels across all birds (Fig. 4B), with test losses ranging from 0.0387 to 0.1467 (mean: 0.1065 ± 0.0441). These results indicate that the network makes accurate predictions when distinguishing syllable segments from gaps and generalizes well across different individual birds. Early stopping with a patience of 5 epochs prevented overfitting, with training terminating after 12–24 epochs (mean: 17 ± 5 epochs; Table S1).

**Figure 4.**
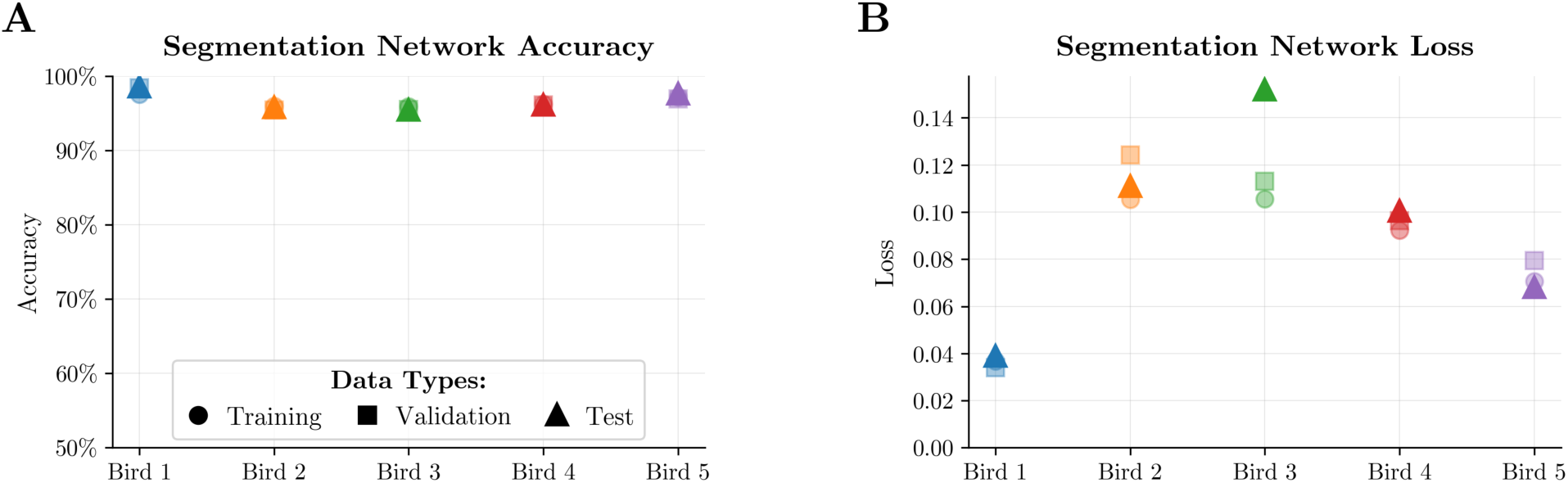
Segmentation network performance. Separate networks were trained for each bird on manually verified song bouts. (A) Segmentation network accuracy across training (circles), validation (squares), and test (triangles) datasets for each of the five birds. (B) Segmentation network loss across training (circles), validation (squares), and test (triangles) datasets for the same five birds.

For example bird 1, which was used in the learning experiment, training stopped at epoch 18 with the best model saved at epoch 13, achieving 98.50% test accuracy. To assess generalization across different data splits, we performed 5-fold cross-validation on bird 1’s training data, which yielded consistent performance across all folds (mean accuracy: 98.52% ± 0.07% standard deviation). The network’s inference speed was approximately 0.73 ms ± 0.1 ms per audio chunk (batch size = 1) on a standard CPU, enabling real-time processing given that new 64-sample chunks arrive every 1.45 ms at the 44.1 kHz sampling rate.

When evaluated for bird 1 against manually verified syllable boundaries with a tolerance of ±15 ms, the sliding window algorithm (see Methods) achieved 96.87% precision (proportion of detected syllable segments that were correct), 98.73% recall (proportion of true syllable segments that were detected), and an F1 score of 97.79%. These results demonstrate that the segmentation network provides accurate syllable boundary detection suitable for real-time applications.

### Classification performance

Classification networks were trained for each of the five birds to assign syllable labels based on spectrograms computed from the first 30 ms after syllable onset (Fig. 5). Training used the same manually verified bouts as for segmentation, comprising 108 to 576 labeled syllable segments across birds (mean: 234 ± 195), which were randomly split into training (70%), validation (15%), and test (15%) subsets (see Table S2 for individual bird data). Across all birds, training and validation accuracies converged to high levels (Fig. 5A), with test accuracies ranging from 96.12% to 97.84% (mean: 97.17 ± 0.82%). Training and validation losses decreased to low levels (Fig. 5B), with test losses ranging from 0.0700 to 0.2057 (mean: 0.1254 ± 0.0544). Early stopping with a patience of 5 epochs prevented overfitting, with training terminating after 10–17 epochs (mean: 14 ± 2 epochs; Table S2). Across birds classification performance was consistent, indicating robust generalization across individual syllable repertoires. For example, bird 1, training stopped at epoch 15 with the best model saved at epoch 10, achieving 97.69% test accuracy. Five-fold cross-validation on bird 1’s training data confirmed consistent performance across all folds (mean accuracy: 97.51% ± 0.70%). The network’s inference speed was approximately 0.94 ms ± 0.25 ms per spectrogram, enabling real-time classification in a separate processing thread.

**Figure 5.**
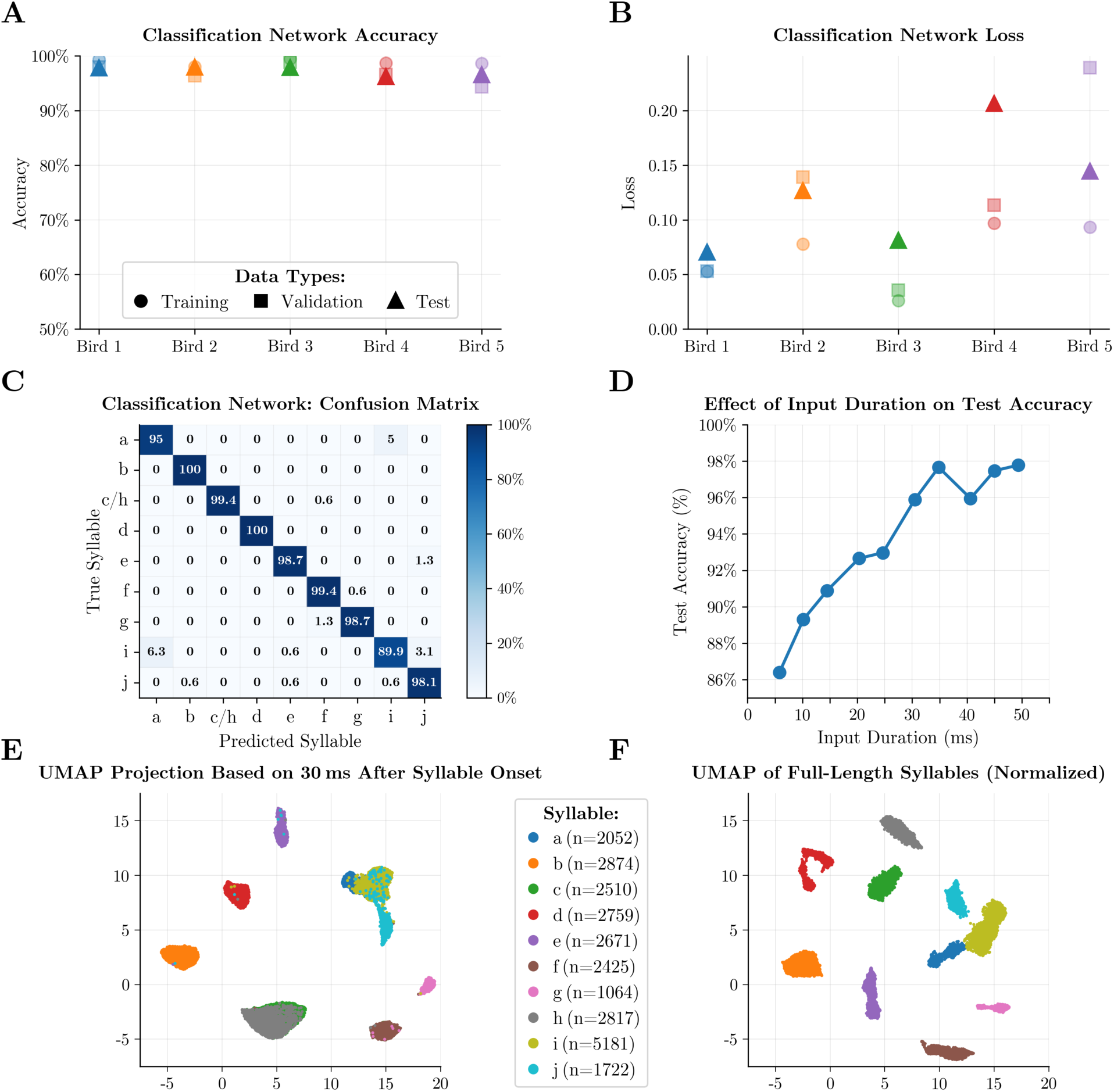
Classification network performance. Networks were trained on syllable spectrograms from the first 30 ms after onset. (A) Classification accuracy for training (circles), validation (squares), and test (triangles) datasets across five birds. (B) Classification loss for training (circles), validation (squares), and test (triangles) datasets across five birds. (C) Confusion matrix for bird 1 (test set, 9 classes with c/h merged). Numbers indicate percentage of syllables classified as each type. (D) Effect of input duration on test accuracy for bird 1 (10 classes, no merging) for input windows from 5-50 ms after syllable onset. (E) UMAP projection of spectrograms using only the first 30 ms after syllable onset (bird 1). Each point represents one syllable, colored by type. N indicates sample sizes for each syllable type. (F) UMAP projection of full-length spectrograms (normalized duration) for the same bird and same syllables as in E.

For bird 1, we examined classification performance in detail using a confusion matrix (Fig. 5C). We chose to merge syllables ‘c’ and ‘h’ for the learning experiment because they were spectrally similar within the first 30 ms and were not targets for online recognition. This allowed us to improve syllable classification accuracy on the remaining syllables without affecting the outcome of the learning experiment. The confusion matrix shows that most syllable types were classified with >98% accuracy. Lower accuracies occurred for syllables with similar spectral structure in the first 30 ms: syllable ‘i’ tended to be confused with ‘a’ (6.3%) and ‘j’ (3.1%), while ‘a’ was misclassified as ‘j’ in 5% of the cases. These confusions reflect genuine spectral similarity in the early portions of these syllables, as visualized in the UMAP projection based on the first 30 ms (Fig. 5E; clusters for ‘i’, ‘j’, and ‘a’ show substantial overlap). In contrast, when using full-length syllables, all syllable types form well-separated clusters (Fig. 5F), confirming that the observed classification errors stem from limited temporal information rather than inherent ambiguity.

To assess the trade-off between classification accuracy and latency, we tested classification performance across input durations from approximately 5–50 ms for the same bird (Fig. 5D). Accuracy increased steeply up to 30 ms (reaching 95.89% without the merging of labels ‘c’ and ‘h’) and showed only modest improvements thereafter. Even at very short input durations (5.8 ms; corresponding to four input chunks), the network achieved 86.39% accuracy, demonstrating substantial discriminability from syllable onsets alone. Based on these results, we selected 30 ms as the default input window for Moove, balancing high accuracy with low latency for real-time experiments. This parameter can be adjusted depending on the specific experimental requirements and song structure.

### Latency characterization

A critical requirement for effective operant conditioning is low-latency feedback delivery. To characterize Moove*’*s latency performance, we measured the time interval between syllable onset detection and white noise (WN) playback onset across different audio buffer configurations. Using bird 2 with networks trained on its individual song repertoire, we targeted a fixed syllable sequence and measured latency for each triggered event. As shown in Figure 6A, latency was quantified as the time from the detected syllable onset (green vertical dashed line) to the onset of WN playback.

**Figure 6.**
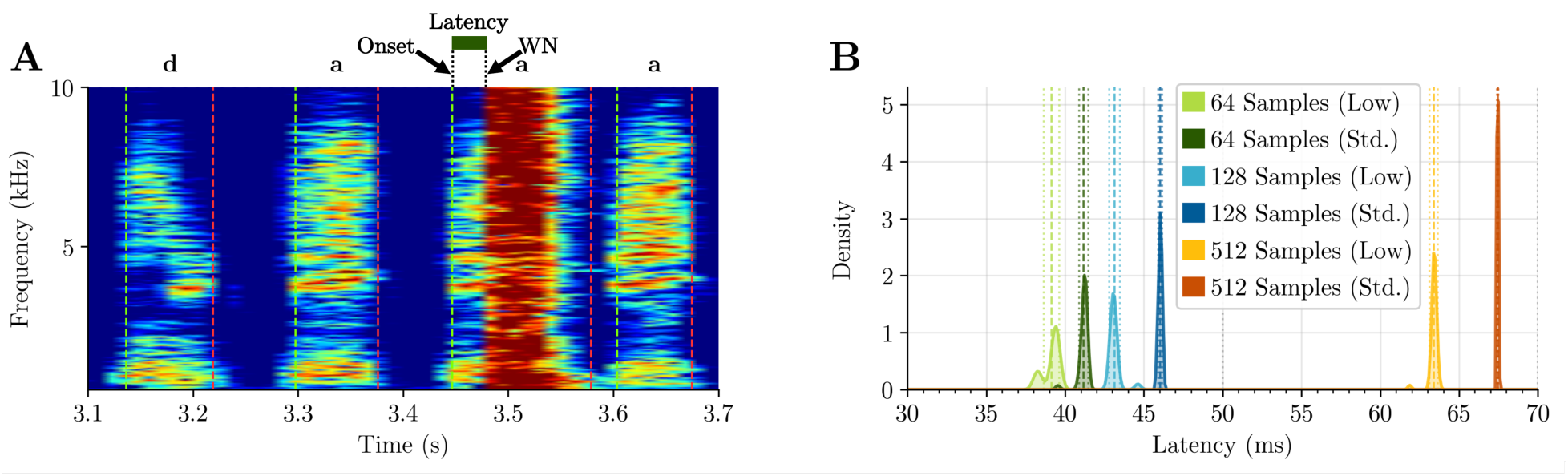
Latency characterization across audio buffer configurations. (A) Example spectrogram showing latency measurement. Green vertical dashed lines mark detected syllable onsets. Latency (green bar) is the time interval between syllable onset and white noise (WN) onset. (B) Distribution of measured latencies for six audio buffer configurations. Three buffer sizes (64, 128, 512 samples) tested with two driver modes (Low Latency or Standard) using the Yamaha Steinberg USB Driver on Windows. Each distribution represents ≥ 50 triggered events.

For the audio interface in our setup, we used the Yamaha Steinberg USB Driver on Windows and tested the *‘*Low Latency*’* and *‘*Standard*’* driver modes with adjustable buffer sizes. Larger buffer sizes provide more stable audio playback by reducing the risk of buffer underruns (audio dropouts), but increase latency. We systematically tested playback latency with three buffer sizes (64, 128, and 512 samples) with the two driver modes (Fig. 6B). Buffer size had a substantial effect on latency: 64-sample buffers produced the shortest latencies (mean: 39.14 ± 0.49 ms in Low Latency mode, 41.18 ± 0.30 ms in Standard mode), while 512-sample buffers resulted in longer latencies (mean: 63.39 ± 0.26 ms in Low Latency mode, 67.45 ± 0.05 ms in Standard mode). Driver mode also affected latency, with Low Latency mode reducing mean latency by approximately 6.01 ± 11.64 ms compared to Standard mode across all buffer sizes. For each configuration, we collected at least 50 triggered instances to obtain reliable latency distributions (n = 60-231 per condition).

### Low-latency feedback performance of Moove

With optimized settings (64-sample buffer, Low Latency mode), mean feedback latencies were 39.14 ± 0.49 ms. Given that Moove waits 30 ms after syllable onset to gather acoustic information for classification, the remaining system overhead (hardware, audio processing, playback) accounts for only ∼10 ms of the latencies. This low-latency performance is suitable for operant conditioning experiments where temporal contingency between behavior and feedback is critical.

### Repeat targeting with position-specific feedback

To study behavioral influences on syllable repetition, we were particularly interested in targeting individual instances of the same syllable within repeat phrases. For example, we might want to target only the fifth occurrence of syllable *‘*a*’* in a phrase consisting of repetitions of syllable *‘*a*’* (Fig. 7). Repeat phrases can be particularly challenging for template-based approaches. They require setting parameters for repetitions of template matches, with refractory periods after matches ensuring that there can be no other matches within the same syllable. Due to subtle changes in the timing of syllables and gaps throughout the repeat phrase, it can be challenging or impossible to define templates and refractory periods in a way that they do not become misaligned over the duration of the phrase, eventually leading to counting errors. Moove*’*s syllable-based approach eliminates this source of counting errors and ensures that latency remains stable with respect to syllable onsets throughout the phrase. As in other sequences, the target can be chosen through pattern matching of regular expressions, allowing the user to specify which position(s) to target within the repeat phrase. As Moove explicitly detects syllable onsets and saves the history of previously detected syllables, it eliminates the need to discriminate spectrally similar syllables based on timing alone. To demonstrate this capability, we show an example song bout in which we targeted the fifth repetition of syllable *‘*a*’* within a repeat phrase for auditory feedback manipulations (Fig. 7). In both phrases in the example bout, Moove successfully detected the fourth *‘*a*’* (a_4_) and triggered playback of different auditory stimuli (*‘*c*’* and *‘*d*’*; red bars). Playback latency was chosen so that playback overlaps with the fifth *‘*a*’* (a_5_). This demonstrates that Moove allows precise control over the target position within repeat phrases.

**Figure 7.**
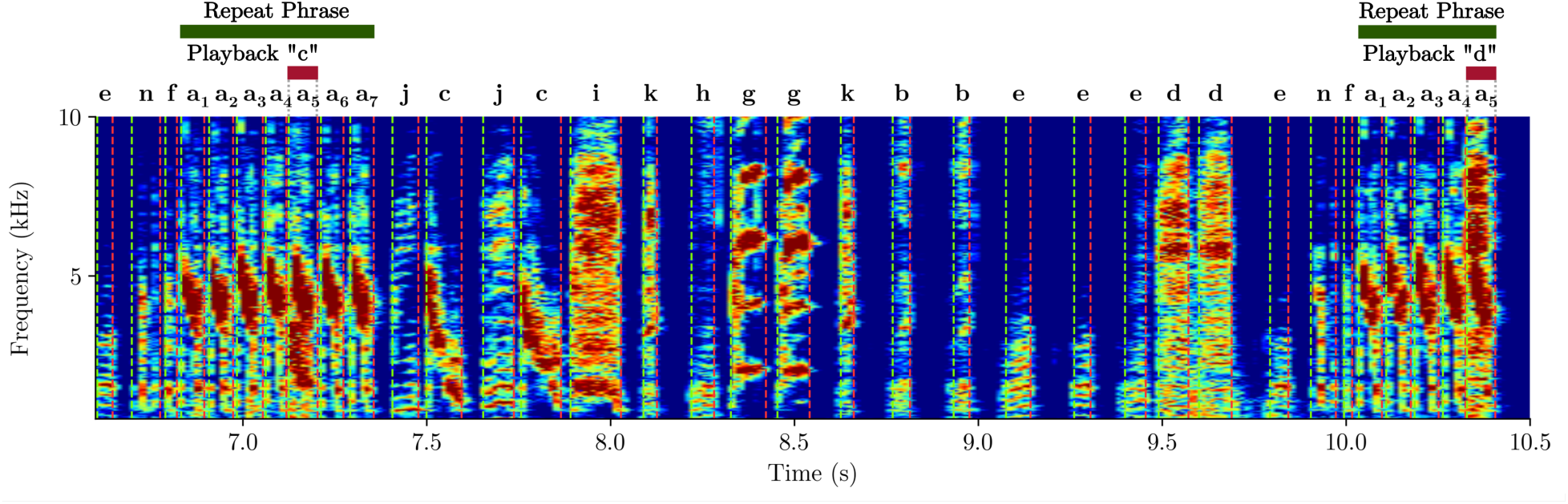
Repeat targeting with position-specific auditory feedback. Example bout demonstrating targeting of the fourth syllable in a repeat phrase of syllable ‘a’ in bird 3. Green bars indicate repeat phrase ‘a’. Red bars indicate auditory playback of different auditory stimuli (‘c’ and ‘d’). Vertical dashed lines mark detected syllable onsets. Latency was set so that playback overlaps syllable a_5_.

### Sequence modification learning experiment

To test whether Moove allows effective operant conditioning, we conducted a sequence modification learning experiment in one bird. This bird exhibited a branch point following the syllable sequence ‘h-c’ during baseline recordings (Fig. 8A), singing either ‘h-c-a’ in 65% or ‘h-c-b’ in 32% of the instances, averaged across the baseline days (Fig. 8B). We chose to target syllable sequence ‘h-c-b’ with white noise feedback, which typically leads to a reduction of the target sequence over multiple training days (Fortkord & Veit, 2025; Veit et al., 2021; Warren et al., 2012). The example spectrogram (Fig. 8A) illustrates the selective targeting of one branch: the sequence ‘h-c-a’ (blue bar) was correctly not targeted, while ‘h-c-b’ (red bar) was accurately detected and the rest of syllable ‘b’ covered by a white noise (WN) playback stimulus. To disentangle learning-related changes from acute effects of WN on subsequent song, 10% of detected sequences were designated as catch trials where no WN was played despite detection of the target sequence. Note that this is different from the typical definition of catch trials and has subsequently been changed in Moove.

**Figure 8.**
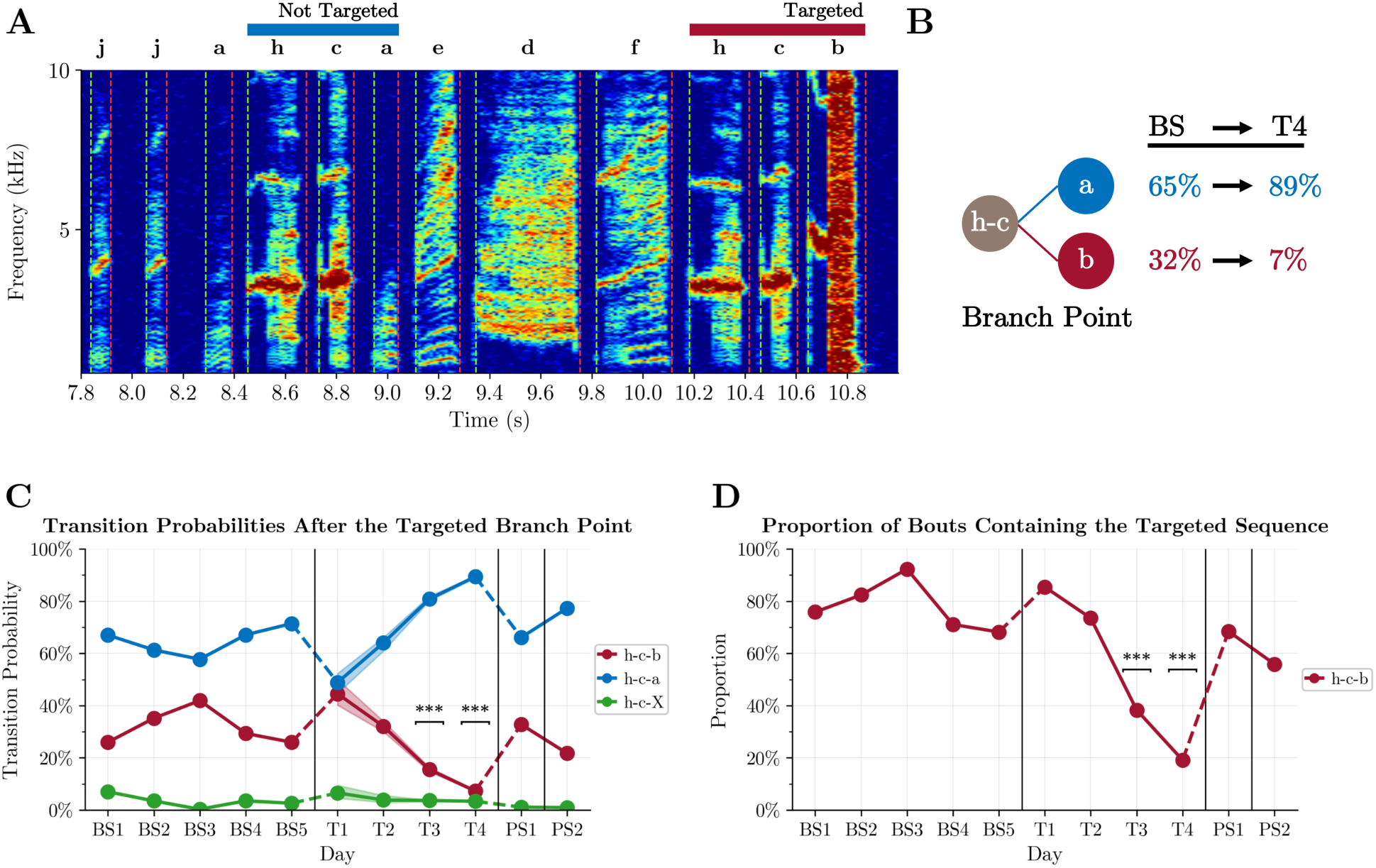
Sequence modification learning experiment using white noise feedback. (A) Example spectrogram from a training day showing selective targeting. Syllable labels indicate detected types online. Left: sequence ‘h-c-a’ (blue bar), not targeted. Right: sequence ‘h-c-b’ (red bar) followed by white noise playback over syllable ‘b’. Vertical dashed lines mark detected syllable onsets and offsets. (B) Summary diagram of learning outcome. Left: mean baseline (BS1–BS5) transition probabilities from ‘h-c’ to ‘a’ (blue, 65%) and ‘b’ (red, 32%). Right: After training (T4), probabilities shifted to 89% for ‘a’ and 7% for ‘b’. (C) Transition probabilities across experimental days. Red: ‘h-c-b’ (targeted, median values); blue: ‘h-c-a’ (main alternative branch); green: other rare transitions. Shaded regions show min-max range from catch-trial adjusted resampling (1000 iterations). Vertical lines indicate non-continuous days between baseline (BS1–BS5), training (T1–T4), and post-screening (PS1– PS2) periods. Statistical significance: *** p < 10⁻¹² (two-proportion z-test). (D) Proportion of bouts containing the target sequence ‘h-c-b’ across days. Statistical significance as in C.

Training with WN feedback resulted in clear behavioral changes. By the final training day (T4), transition probabilities at the target branch point had shifted from the baseline average: transitions to ‘a’ increased to 89% while transitions to ‘b’ decreased to 7% (Fig. 8B), demonstrating successful sequence modification which reduced the number of WN feedbacks. Figure 8C shows the full experimental timeline from baseline across all training days and post-screening days. In line with the steady reduction in the target branch over the training days (T1-T4), the bird increased the alternative branch ‘a’ after ‘h-c’ (blue line). In this experiment, catch-trials were only assigned to bouts containing the target sequence. To account for bias from this incomplete sampling, we performed a resampling procedure: we randomly sampled 10% of all bouts and therein replaced any non-catch-trial bouts containing the target sequence with randomly sampled catch-trial bouts containing the target sequence (with replacement), repeating this 1000 times. This simulates the effect that the missing catch trial information can have on the analysis. From the resulting distribution, we extracted minimum, median, and maximum transition probabilities to quantify uncertainty. Shaded regions in Figure 8C show the range between minimum and maximum values from this resampling. Statistical analysis using two-proportion z-tests on both median and maximum values (representing the most conservative estimate) confirmed that from training day T3 onward, the reduction in ‘h-c-b’ transitions was highly significant compared to baseline (median: T3 p = 3.24 × 10⁻²⁰, T4 p = 1.73 × 10⁻¹²; maximum: T3 p = 3.17 × 10⁻¹⁸, T4 p = 1.73 × 10⁻¹²). The proportion of bouts containing the targeted sequence ‘h-c-b’ also decreased dramatically, from 70–85% during baseline to 19% on day T4 (Fig. 8D), with highly significant reductions from day T3 onward (T3: p = 1.87 × 10⁻¹⁴; T4: p = 1.78 × 10⁻²⁴). In post-screening sessions conducted after training, transition probabilities and target sequence occurrence had partially recovered toward baseline (PS1 vs baseline: p = 1.08 × 10⁻⁵).

Throughout the experiment, Moove performed reliably: zero false-positive WN triggers occurred across all four training days. False negatives (missed triggers) were <1%. In this experiment (performed before the optimization of audio driver parameters reported in Fig. 6), mean WN onset latency measured from onset of syllable ‘b’ was 90.87 ms ± 11.04 ms (n = 315 triggers). These results demonstrate that Moove enables accurate, low-latency syllable targeting suitable for reinforcement learning experiments, with behavioral outcomes comparable to those achieved using template-based real-time systems.

## Discussion

We developed Moove, a two-stage neural network system that enables real-time syllable targeting during ongoing Bengalese finch song. The name is derived from “Japanische Mövchen”, the German name for Bengalese finches. Across multiple birds, Moove achieved high segmentation and classification accuracy with low feedback latencies. A sequence modification learning experiment validated the system’s effectiveness for closed-loop operant conditioning. Moove achieves feedback delivery before syllable offset, a capability previously limited to template-based systems, while offering advantages in syllable coverage, classification accuracy, and experimental flexibility.

### Moove can target ongoing syllables

The central innovation of Moove is its ability to classify and target syllables using neural networks before their acoustic offset, addressing a critical limitation in the current toolset for closed-loop experiments. Existing real-time neural network systems for birdsong classification (Kawaji et al., 2024; Schulthess et al., 2023; Steinfath et al., 2021) determine syllable identity only after detecting syllable offset. While this approach is suitable for continuous monitoring or offline analysis, it precludes feedback delivery during ongoing vocalizations. This is critical for many operant conditioning experiments, where temporal contingency between behavior and outcome affects learning efficacy (Charlesworth et al., 2011; Tumer & Brainard, 2007). For many experimental protocols, including auditory feedback manipulation, song-triggered optogenetic stimulation, or reinforcement learning protocols, the ability to deliver feedback while the bird is still vocalizing the target syllable is essential.

TweetyNet (Cohen et al., 2022) is the state-of-the-art for offline syllable annotation, combining convolutional feature extraction with bidirectional LSTM layers to capture temporal dependencies across large spectrogram windows. This architecture achieves excellent accuracy by processing extensive temporal context both before and after each time point, making the current architecture incompatible with real-time targeting. TinyBird-ML (Schulthess et al., 2023) employs a two-stage architecture similar to Moove, but is designed for deployment on wearable sensors worn on birds’ backs for real-time classification in social groups. The wearable design imposes constraints including battery limitations and limited temporal resolution for power conservation. SAIBS (Kawaji et al., 2024) uses a two-stage approach with threshold-based segmentation that relies on very long temporal windows. These windows likely include information from preceding syllables, presumably creating position-dependent subclusters in which the same syllable type receives different labels depending on sequence context. The Deep Audio Segmenter (DAS; Steinfath et al., 2021) represents the most versatile existing tool, using temporal convolutional networks to handle diverse vocalizations from insect sounds to complex birdsong. Like Moove, DAS includes a GUI for dataset creation and training. Like other existing neural network-based tools, DAS assigns syllable identity only after offset detection.

To achieve pre-offset classification, we prioritize temporal precision over marginal accuracy gains from extensive temporal context or bidirectional processing. Moove therefore employs a forward-only architecture with a small temporal window, enabling syllable classification during ongoing vocalizations while maintaining accuracy suitable for real-time targeting. With this design choice, Moove fills the gap between offline neural network-based tools and template-based real-time systems.

### Moove classifies syllables using only local acoustic information

Syllables in Moove are classified based solely on their own acoustic properties, without incorporating information about preceding syllables. While incorporating sequential context could dramatically improve accuracy by exploiting probabilistic sequence structure, this would introduce systematic bias against detecting rare or novel syllable transitions. For experiments investigating vocal sequencing or designed to modify sequences through operant conditioning, context-independent classification is essential. A context-dependent classifier would become progressively less accurate as birds learn to avoid targeted sequences, requiring retraining during experiments or obscuring sequence modifications through mislabels based on sequential expectations of the classifier.

### Moove performs syllable-based classification and can target syllables based on sequence context

Template-based systems like EvTAF (Ali et al., 2013; Moll et al., 2023; Riedner & Adam, 2020; Roberts et al., 2012; Trusel et al., 2025; Tumer & Brainard, 2007; Xiao et al., 2018) recognize syllables by matching a short audio segment to a spectral template, achieving very low latencies when combined with specialized hardware. However, template creation is labor-intensive, requiring manual design and iterative optimization for each target syllable. Even with optimization algorithms, only 18–43% of syllables in Bengalese finch repertoires can be detected reliably using manually created templates (Kulkarni & Troyer, 2020), which fundamentally limits experimental accessibility. When birds modify their syllables during learning experiments, fixed templates become less effective and require manual updating.

Defining context-dependent targets compounds these challenges, requiring complex combinations of templates and timing constraints, with logic operators that can become unwieldy as sequence complexity increases. Because Moove explicitly identifies each syllable as it occurs, experimenters can easily target syllables based on their identity and the identity of preceding syllables (‘sequence context’). This syllable-based approach proves particularly valuable for Bengalese finch song, where acoustically identical syllables can appear in multiple sequence contexts (such as ‘d-e1-f’, or ‘x-e2-y’) (Jin & Kozhevnikov, 2011; Katahira et al., 2011; Koparkar et al., 2024; Lu et al., 2025; Veit et al., 2021). Experimenters can specify even complex target sequences by listing the syllable labels with regular expression–based pattern matching. A template-based approach is particularly difficult for repeat phrases, where subtle changes in syllable and gap timing throughout the phrase cause template-timing combinations to become misaligned, eventually leading to counting errors. Moove’s syllable-based approach is intuitive to use and accommodates natural variability in song structure and timing. It eliminates timing-based counting errors in repeat phrases and ensures stable feedback latencies relative to syllable onsets (Fig. 7). Syllable-based detection also reduces false-positive triggers from call vocalizations or cage noise, which can spuriously match acoustic templates.

### Moove does not require specialized hardware or skills

Moove’s Python-based design facilitates modification and extension, enabling researchers to adapt the system to specific experimental needs. Moove’s source code is released under the MIT License. The system operates on standard desktop recording hardware with commodity audio interfaces. File formats and annotation structures maintain compatibility with existing workflows using EvTAF, facilitating adoption by research groups transitioning from these systems. MooveGUI provides a complete pipeline from data collection through network training, where researchers can train bird-specific models without deep machine learning expertise. The semi-automated approach integrates algorithmic segmentation and unsupervised clustering with manual tools for correction and refining cluster memberships. The modular architecture permits researchers to modify segmentation and classification components separately for specific experimental needs.

### Moove is suitable for reinforcement learning experiments

The sequence modification learning experiment validated Moove’s effectiveness for closed-loop operant conditioning. After targeting one branch of a branch point with white noise feedback, the bird exhibited a clear shift in transition probabilities from baseline. These learned changes are comparable to published sequence learning experiments (Veit et al., 2021; Warren et al., 2012; Fortkord & Veit, 2025). In this experiment, mean white noise onset latency was approximately 90 ms, as it was conducted before the audio driver optimizations that reduced latency to approximately 45 ms in later testing. Despite this higher latency, the bird learned effectively, suggesting that this temporal contingency can be sufficient for operant conditioning in a sequence learning task. Our pilot results demonstrate that Moove enables accurate, reliable syllable targeting suitable for reinforcement learning experiments, with behavioral outcomes comparable to those achieved using template-based systems.

### Limitations and Future Perspectives

Several aspects of Moove’s current implementation present opportunities for extension. The feedback latency of approximately 45 ms proved sufficient for sequence learning but may require reduction for pitch learning or other experiments (Tumer & Brainard, 2007). As the latency is composed of the 30 ms default classification window and 15 ms system overhead, this could be achieved through shorter classification windows, at the cost of accuracy. Shortening the adjustable window to approximately 15 ms would make classification performance and latency comparable to EvTAF. This trade-off is user-adjustable without code modification. Hardware optimizations or more complex network architectures could potentially improve discrimination without extending the temporal window. If target syllables occur in a fixed sequence, the classification window could be limited to the window necessary for pitch calculation.

The segmentation network occasionally misidentifies environmental noise as syllable segments, though this did not affect targeting in our experiment. Future implementations could include a dedicated noise class during training, as implemented in other systems (Cohen et al., 2022; Steinfath et al., 2021).

Applicability of Moove to vocal signals in other species will depend on their vocal structure (Sainburg et al., 2020). Songbird species with discrete syllable repertoires (e.g., zebra finches) should be well-captured by Moove’s approach. Bird species with highly variable or continuous vocalizations (Cohen et al., 2020; Costalunga et al., 2023; Markowitz et al., 2013; Zhao et al., 2023) present greater challenges due to less discrete syllable boundaries and may require substantial modifications to segmentation and classification approaches. Extension to closed-loop experiments requiring the detection of vocalizations in other species (Hage et al., 2013; Liao et al., 2024; Pomberger et al., 2018) may be easily achieved, potentially after adaptations to different acoustic ranges and temporal scales.

Moove’s real-time targeting capability supports diverse experimental applications beyond the operant conditioning protocol presented here. The system could be used, for example, for song-triggered manipulations of auditory feedback (Tumer & Brainard, 2007), visual feedback (Zai et al., 2020), neural microstimulation (Moll et al., 2023), optogenetic perturbations (Hisey et al., 2018), and access to conspecifics (Ben-Tov et al., 2023). The open-source release will enable community-driven extensions and validation across diverse experimental contexts and species. For experimental protocols involving sequence targeting, Moove’s combination of broad syllable coverage, high accuracy, flexible sequence specification, and moderate latency presents a practical solution for previously inaccessible experimental approaches.

## Code availability

The source code for Moove is available at https://github.com/veitlab/moove.

## Acknowledgements

This work was supported by a Daimler Benz Postdoc Fellowship to LV (grant number 31-10/21) and the Deutsche Forschungsgemeinschaft (DFG, German Research Foundation) – Project number 536953998. FH is supported by a PhD fellowship by the Ersatzausschreibung Landesgraduiertenförderung of the Excellence Strategy of the University of Tübingen. We thank members of the Veit lab Avani Koparkar, Lioba Fortkord, Priya Binwal and Abhilipsa Das for help with testing Moove or comments on the manuscript.

## Author contributions

NR and LV designed the study, NR implemented Moove, gathered data, analyzed the data, JLG and FH gathered additional data and contributed to code and analysis, NR and LV wrote the paper, JLG and FH review and editing.

---

**Table S1:** Segmentation network performance across five birds.

| Bird ID | Total Bouts | Total Epochs | Training Accuracy (%) | Validation Accuracy (%) | Test Accuracy (%) | Training Loss | Validation Loss | Test Loss |
| --- | --- | --- | --- | --- | --- | --- | --- | --- |
| Bird 1 | 576 | 18 | 97.58 | 98.45 | 98.50 | 0.0368 | 0.0339 | 0.0387 |
| Bird 2 | 108 | 18 | 94.92 | 95.24 | 94.30 | 0.1323 | 0.1213 | 0.1450 |
| Bird 3 | 113 | 14 | 96.09 | 96.35 | 96.26 | 0.0982 | 0.0926 | 0.0974 |
| Bird 4 | 176 | 24 | 95.08 | 95.32 | 94.46 | 0.1299 | 0.1210 | 0.1467 |
| Bird 5 | 199 | 12 | 95.84 | 95.79 | 96.04 | 0.1115 | 0.1150 | 0.1048 |
| Mean ± SD | 234 ± 195 | 17 ± 5 | 95.90 ± 1.06 | 96.23 ± 1.32 | 95.91 ± 1.70 | 0.1017 ± 0.0389 | 0.0968 ± 0.0371 | 0.1065 ± 0.0441 |

**Table S2:** Classification network performance across five birds.

| Bird ID | Total Bouts | Syllable Classes | Total Epochs | Training Accuracy (%) | Validation Accuracy (%) | Test Accuracy (%) | Training Loss | Validation Loss | Test Loss |
| --- | --- | --- | --- | --- | --- | --- | --- | --- | --- |
| Bird 1 | 576 | 9 | 15 | 99.34 | 97.96 | 97.69 | 0.0529 | 0.0530 | 0.0700 |
| Bird 2 | 108 | 8 | 14 | 98.03 | 96.43 | 97.84 | 0.0778 | 0.1389 | 0.1263 |
| Bird 3 | 113 | 8 | 17 | 99.51 | 98.85 | 97.77 | 0.0261 | 0.0356 | 0.0808 |
| Bird 4 | 176 | 10 | 10 | 98.71 | 96.63 | 96.12 | 0.0968 | 0.1135 | 0.2057 |
| Bird 5 | 199 | 8 | 15 | 98.63 | 94.33 | 96.45 | 0.0933 | 0.2392 | 0.1441 |
| Mean ± SD | 234 ± 195 | 9 ± 1 | 14 ± 2 | 98.84 ± 0.59 | 96.84 ± 1.72 | 97.17 ± 0.82 | 0.0694 ± 0.0297 | 0.1160 ± 0.0808 | 0.1254 ± 0.0544 |

